# Achieving Reversible Ligand-Protein Unbinding with Deep Learning and Molecular Dynamics through RAVE

**DOI:** 10.1101/400002

**Authors:** João Marcelo Lamim Ribeiro, Pratyush Tiwary

**Affiliations:** Department of Chemistry and Biochemistry and Institute for Physical Science and Technology, University of Maryland, College Park 20742, USA

## Abstract

In this work we demonstrate how to leverage our recent iterative deep learning–all atom molecular dynamics (MD) technique “Reweighted autoencoded variational Bayes for enhanced sampling (RAVE)” (Ribeiro, Bravo, Wang, Tiwary, J. Chem. Phys. 149, 072301 (2018)) for sampling protein-ligand unbinding mechanisms and calculating absolute binding affinities when plagued with difficult to sample rare events. RAVE iterates between rounds of MD and deep learning, and unlike other enhanced sampling methods, it stands out in simultaneously learning both a low-dimensional physically interpretable reaction coordinate (RC) and associated free energy. Here, we introduce a simple but powerful extension to RAVE which allows learning a position-dependent RC expressed as a superposition of piecewise linear RCs valid in different metastable states. With this approach, we retain the original physical interpretability of a RAVE-derived RC while making it applicable to a wider range of complex systems. We demonstrate how in its multi-dimensional form introduced here, RAVE can efficiently simulate the unbinding of the tightly bound benzene-lysozyme (L99A variant) complex, in all atom-precision and with minimal use of human intuition except for the choice of a larger dictionary of order parameters. These simulations had a 100 % success rate, and took between 3–50 nanoseconds for a process that takes on an average close to few hundred milliseconds, thereby reflecting a seven order of magnitude acceleration relative to straightforward MD. Furthermore, without any time-dependent biasing, the trajectories display clear back–and– forth movement between various metastable intermediates, demonstrating the reliability of the RC and its probability distribution learnt in RAVE. Our binding free energy is in good agreement with other reported simulation results. We thus believe that RAVE, especially in its multi-dimensional variant introduced here, will be a useful tool for simulating the dissociation process of practical biophysical systems with rare events in an automated manner with minimal use of human intuition.

## 1 INTRODUCTION

Modern all-atom simulation approaches to tackling open problems in the biological sciences must confront an inherent limitation stemming from the difference in timescales between the fast rovibrational molecular motions relative to the (much) slower biochemical processes. With these fast molecular motions acting to constrain the timestep for integrating Newton’s equations of motion to small femtosecond values, it can be difficult to simulate timescales much greater than a few tens of microseconds despite intense developments in computational hardware. ^1^ It is often the case, however, that fundamental biological processes reach timescales greater than milliseconds, one particular example being the (un)binding of specific ligand-protein complexes happening over multiple hours.^2,3^ In order to deal with this restriction several methods have been introduced with the aim of reducing the timescale for sampling the slow biochemical processes while nonetheless being capable of recovering their original statistics. ^4–18^ Included among these are several interesting recent approaches that leverage deep learning to generate an optimum reaction coordinate (RC) that can then be used within a pre-existing enhanced sampling or markov state model framework. ^11–15^ One of these is a method we very recently proposed, named “Reweighted autoencoded variational Bayes for enhanced sampling (RAVE)”. The major distinguishing feature of RAVE is that the RC is learnt together with its Boltzmann probability distribution,^18^ which can then serve as the ideal bias potential and be leveraged outside pre-existing biasing frameworks such as metadynamics or umbrella sampling.^4–6,10^ In the original proof-of-concept paper, RAVE was applied to model potentials including a fullerene-nanopocket ^19–23^ unbinding test case where it was demonstrated that sampling in simulations could indeed be enhanced with the simultaneous on-the-fly learning of the RC and bias potential.^18^ We demonstrated that RAVE could reproduce the dissociation free energy profile for the unbinding of a fullerene from a nanopocket in much less computational time than using the popular metadynamics and umbrella sampling methods.^24^ These initial investigations were thus suggestive that RAVE could find important applications in simulations of ligand-protein complexes that now have a critical role in aiding drug design.^25,26^

Here we introduce a simple but powerful multi-dimensional extension to RAVE that makes it possible to obtain accurate absolute binding free energies Δ*G*_*b*_ in realistic ligand-protein complexes. As our test-case, we choose benzene (un)binding from the L99A variant of the T4 bacteriophage lysozyme protein (T4L), which is a popular and challenging ligand-protein complex for experimental and simulation studies. ^25,27–35^ For example, Deng and Roux investigated the binding affinities of the T4L protein with various aromatic ligands.^25,30^ Miao and co-workers applied their Gaussian Accelerated Molecular Dynamics (GaMD) method on the complex.^31^ Wang et al. ^33^ also looked into the binding free energies but from the perspective of using association and dissociation rate coefficients to calculate the Δ*G*_*b*_. Most recent, several (un)binding paths were simulated in order to make accurate kinetics predictions,^34,35^ these (un)binding paths differing from one another in that different helix-helix distances were modulating the entry and exit of benzene from the binding pocket.

Using our multi-dimensional extension, we will show how RAVE can learn position-dependent RCs which lead to sampling benzene unbinding from the buried binding pocket of the T4L protein in 100% of our short independent MD simulations (that is 20 out of 20 simulations). The first unbinding event in these short independent runs occurred within 3-50 nanoseconds, corresponding to a seven order of magnitude speed-up relative to the actual process expected to take around 100 milliseconds. Furthermore, back and forth movement between the deep initial bound state and intermediate metastable states within the binding pocket was often observed prior to unbinding. This rate of success and speed-up, together with the reversible nature of the sampling within the buried binding pocket and elsewhere inside the protein, are to our knowledge unprecedented in systems that are this complex. In addition, we also show that RAVE can handle the calculation of binding free energies quite well. Multiple runs gave estimates of the binding free energy within reasonable error bars in agreement with other sampling methods. Lastly, using our simulations we can construct well-converged free energy profiles demonstrating the interdependence between ligand movement, protein fluctuation and water movement. It is our expectation that RAVE, especially with its multi-dimensional extension introduced here, will find use in automating the calculation of important quantities which are of practical interest.

## 2 THEORY

### 2.1 RAVE

RAVE has been introduced in detail in the original proof-of-concept publication^18^ and here we summarize its central features. To begin let us assume a molecular system with *N* atoms at temperature *T*, and under some other generic thermodynamic conditions. Our central objective is to sample the system’s Boltzmannweighted probability distribution using allatom MD. To achieve this objective, the first step in RAVE is to launch an unbiased MD simulation from a point in configuration space that often, for ligand-protein unbinding processes, will correspond to the ground state if known or to a metastable state. Unless *T* is high, the simulation will be trapped in this state sampling the fast internal degrees of freedom as well as the a priori unknown RC, both according to their Boltzmann-weighted probabilities. There is useful information contained in this trapped simulation that can be leveraged to enhance fluctuations. The slow degree-of-freedom describing escape from the metastable state, for instance, stands apart as a distinct signal or feature relative to the fast internal oscillations that together amount to background noise. The probability distribution *P* of the simulation data, when projected onto the slow coordinate, provides a natural biasing potential that helps enhance the fluctuations by making motion along the RC more diffusive. Our central motivation in constructing RAVE was that both the slow degree-of-freedom as well as its probability distribution *P* are learnable via the use of certain forms of deep learning.^18^ RAVE will then proceed from this initial unbiased simulation stage in an iterative fashion, each iteration aiming to simultaneously construct, from what will then be a biased simulation, a better biasing potential along a more refined RC. In this sense, the biasing along the approximate RC obtained from deep learning drives forward the enhanced exploration of the system’s configuration space, leading to the generation of new data that we can again analyze with deep learning. The success of this data generation scheme will depend on the RAVE protocol determining an appropriate RC to describe the slow coordinate, but the iterative protocol per construction provides a self-consistent check to help construct the correct solution. One can just screen through the set of available RAVE RCs, including any spurious solutions, since an appropriate RC should lead to systematically greater exploration of configuration space until ergodicity is achieved. It is the method’s inherent new data generation scheme together with its simultaneous learning of both the RC as well as the bias potential that makes RAVE distinct relative to other deep learning based methods.^12–15^

Let us formalize this intuitive description. To start RAVE the user first defines a *k*dimensional vector **s** whose components are *k* order parameters (*s*_1_, *s*_2_, *…, s*_*k*_) expected to have an important role in the process being studied. These are functions of the 3*N* atomic coordinates **x**, *s*_*i*_ = *s*_*i*_(**x**) where *i* = 1, 2, *…, k*, and represent a valid reduced dimensional description. One should think of a good order parameter vector **s** as containing components of a basis set that together can be combined into a RC that describes the slow degree-of-freedom associated with the process. In the case of ligand-protein complexes, which are the subject of this work, the vector components are distances between the ligand center-of-mass and protein residues, protein inter-residue distances and protein residue hydration states.^36,37^ It is standard practice in the enhanced sampling literature to introduce such order parameter descriptions.^16,36–41^ Although the need to pre-select a set of order parameters might seem like a drawback of RAVE and these other methods,^16,38–40^ in this work we will demonstrate how RAVE allows us to expand an initial minimal list of order parameters with a self-consistent test until the set of order parameters required to construct the RC is complete. This is akin to starting with a small basis set in quantum mechanics, and gradually expanding the list if needed.

With the order parameters in hand we can launch a brief unbiased molecular simulation from the ground state configuration. This generates a time-series (**s**^1^, **s**^2^, *…***s**^*n*^) tracking the time evolution of **s**, where *n* is the total number of timesteps in the simulation, and **s**^*i*^ denotes values of the order parameter vector at time-step *i*. This time-series dataset can then be used to extract, *at the same time*, both (a) a latent variable *z* that describes a low-dimensional manifold capturing the interesting features of the data and (b) probability distribution *P* (*z*) of the data along this variable *z*. Since for our purposes the task of capturing the interesting features amounts to finding a low-dimensional representation of the data that disentangles the slow molecular motions from the rovibrational oscillations amounting to random noise, RAVE uses an unsupervised machine learning approach called variational autoencoder (VAE)^42–44^ whose training protocol per construction compresses high-dimensional data into low-dimensional representations. A crucial point worth noting is that in traditional implementations of the VAE, *z* is described as tens of thousands of neural network parameters lacking clear interpretability. RAVE attempts to solve the interpretability problem by shifting the emphasis to the probability distribution *P* (*z*) of the latent variable, rather than the exact variable itself. By screening through trial RCs *χ* expressed as linear combination of the order parameters (*s*_1_, *s*_2_, *…, s*_*k*_), RAVE determines the best RC as the *χ* whose probability distribution *P* (*χ*) is closest to the probability distribution learnt in VAE, namely *P* (*z*), according to the Kullback-Leibler (KL) divergence metric:

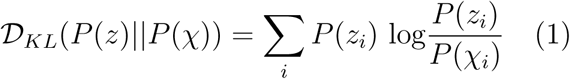

Eq. (1) sums over the discretized distributions containing the same number of bins. In the limit when *P* (*χ*) approaches *P* (*z*), the KL divergence will tend to zero. The distribution *P* (*χ*) minimizing Eq. (1), in addition to determining the RC *χ*, immediately leads to a bias potential, *V*_*b*_, equal to the inverted free energy,

*F* :

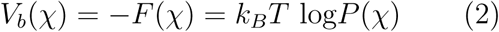

At this point RAVE has determined, from an unbiased simulation, both an approximate reaction coordinate as well as a bias potential, which it uses to launch a short biased MD simulation generating a new dataset (**s**^1^, **s**^2^, *…***s**^*n*^) containing larger fluctuations in configuration space, assuming that the RC identified indeed has sufficient overlap with the true RC. From this new dataset we can then extract a more accurate (in terms of sampling the tails) distribution *P* along a refined hidden latent variable *z* whose unbiased probability distribution we compare to that of trial RCs *χ* again expressed as a linear combination of order parameters (*s*_1_, *s*_2_, *…, s*_*k*_). In order to reweight out the effect of the biasing, we calculate the KL Divergence as:

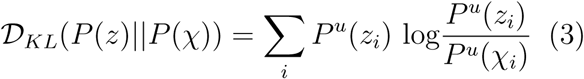

where *P*^*u*^(*z*) and *P*^*u*^(*χ*) are the unbiased probabilities reweighted from a biased MD simulation. For this RAVE associates a weight 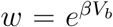 to each point sampled during the biased simulation, where 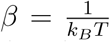 is the inverse temperature. This can then be used to recover the unbiased probabilities through the simple reweighting formula from importance sampling:

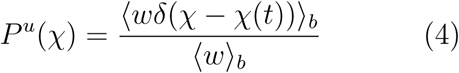

In Eq. 4 the subscript *b* indicates averaging while sampling from the biased simulation. From this point onwards all that is left is to launch another molecular simulation using the current biasing parameters and iterate between rounds of MD and VAE until the desired thermodynamic variables as well as the RC are converged.

### 2.2 Multi-Dimensional RAVE and the “washing out” trick

The original RAVE protocol was designed to determine a one-dimensional RC given a set of user-defined input order parameters.^18^ Furthermore, the one-dimensional RC was restricted to be a linear combination of these order parameters, as described in Sec. 2.1. It can be impractical, however, to use just a single linear RC for problems such as ligand-protein unbinding, where the reaction path involves movements between multiple metastable states. Of course, in principle, we could extend the space of available RCs to include non-linear combinations of the chosen order parameters. These nonlinear RCs together with the associated bias could handle realistic biochemical problems of arbitrary complexity. Unfortunately, the use of non-linear functions is not without their own complications. For instance, when using Eqs. (1) and (3), different non-linear combinations can lead to almost indiscernible values of KL divergence, increasing the likelihood of finding spurious solutions for the optimum RC and bias. Furthermore, the natural limit of taking non-linear combinations of order parameters is to directly use the deep neural network VAE bottleneck variable itself as the RC, similar to what has been done in recent autoencoder based work.^12,13,45^ However, this at odds with our key intention of maintaining physical interpretability of the RC.

Here we develop an alternative approach based on representing the RC as a combination of multiple position-dependent piecewise linear functions. Although the true RC is not embedded in a linear subspace spanned from the user-defined input order parameters, we divide the sampled regions of configuration space into several sections such that within each section the local component of the RC is embedded in a linear subspace of the order parameters. Each linear RC component and its associated Boltzmann probability distribution can then be identified using the original RAVE protocol in one of two different procedures. These can be either (a) in a supervised piecewise manner where the network is trained only using data from specific parts of the configuration space, or (b) in a relatively unsupervised manner where the full dataset can be used, but with a slight modification that we introduce next. This allows enforcing that each local linear component is optimized strictly from only the local features of the configuration space. We adopt a minimalistic approach here, where each subsequent local component is introduced only when no further enhancement in ergodicity can be achieved with the components at hand. Notice that each local component that is introduced is free to learn a different direction in the configuration space of our molecular system such that biases along them act about multiple and ideally independent dimensions. It is for this reason that we call this extension of the protocol multidimensional RAVE (multi-RAVE). We provide a flowchart depiction of multi-RAVE in Fig. 1.

**Figure 1:**
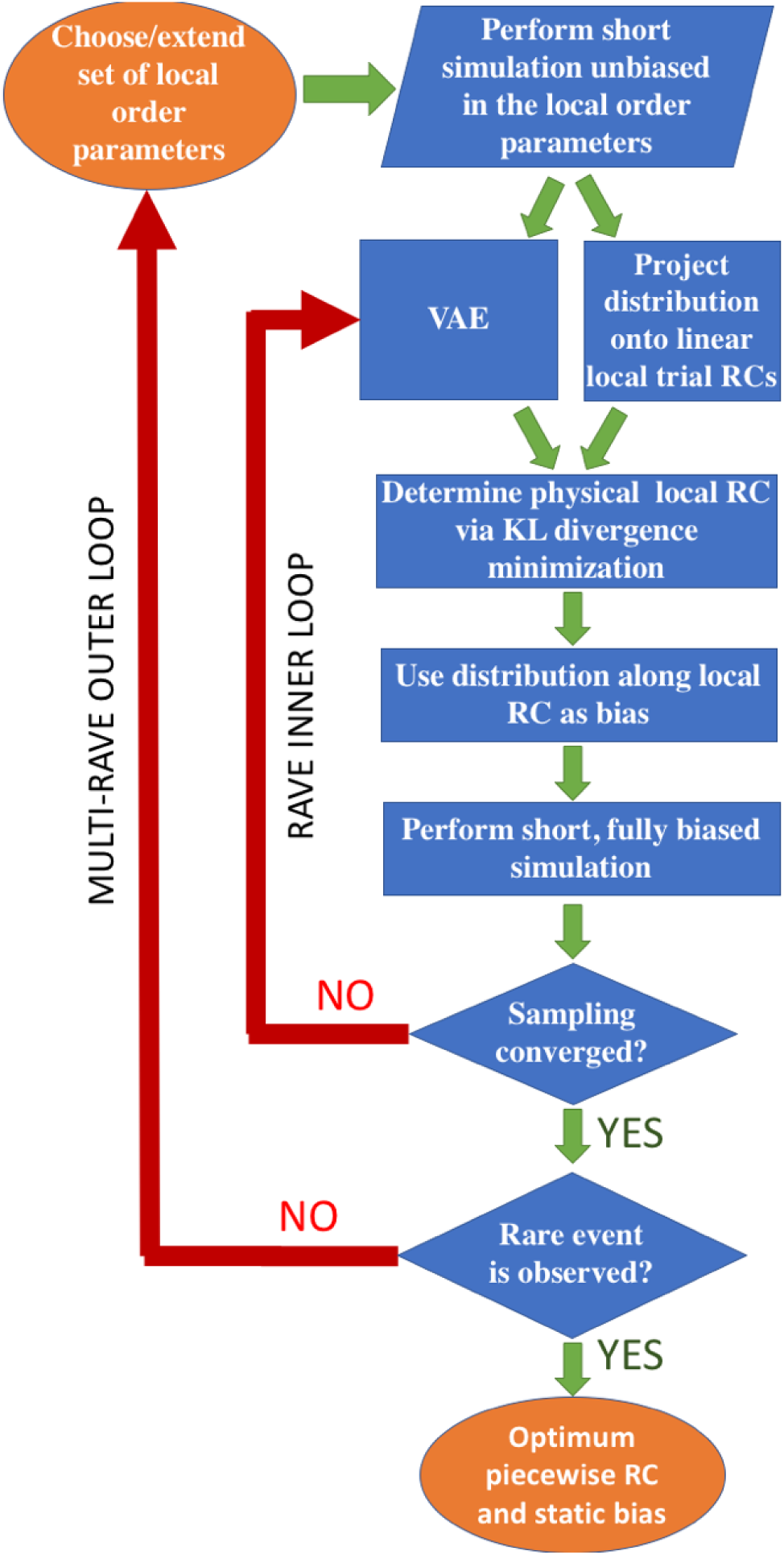
Flowchart illustrating the multi-RAVE protocol. The portions of the flowchart in blue represent the original RAVE protocol while the outermost loop represents the multidimensional extension introduced in this work.

We now formalize this intuitive picture. To initiate multi-RAVE we first launch a short unbiased molecular simulation from an initial configuration that corresponds to the ground state configuration. Using the dataset containing the time evolution of **s** we now subject the simulation data to the original RAVE protocol under the specific constraint that the first RC component, which we call *χ*_1_, be a linear combination of the user-input order parameters **s**, i.e. *χ*_1_ = **c s** = *c*_1_*s*_1_ + *c*_2_*s*_2_ + *…* + *c*_*k*_*s*_*k*_. Nothing until now has distinguished multi-RAVE from the original protocol, since in essence we are performing RAVE on a single linear reaction coordinate. As described in Sec. 2.1 and in the original publication,^18^ RAVE will proceed to iterate between rounds of MD and VAE in order to improve the choice of the coefficients **c** and the associated unbiased probability distribution. Should the optimized one-dimensional RC *χ*_1_ lead to sampling the desired rare event, then nothing else remains to be done. If, however, after initial enhancement in ergodicity the coefficients converge but the system still does not exhibit the desired rare event we set out to study, we proceed to introduce a new round of RAVE on potentially new order parameters.

Essentially, the first set of RAVE has brought the system to an intermediate where either (a) new order parameters are needed, and/or (b) previous order parameters suffice, but the RC undergoes a change which cannot be captured by the constraints of a linear formalism, which is the case with the benzene-T4L complex we will focus on. See Figs. 2b and 2c for an illustrative schematic of case (b). To tackle this issue, multi-RAVE now introduces a second linear RC component, *χ*_2_, that will be optimized within a region of configuration space not sampled during biased MD simulations with RC *χ*_1_ and associated bias *V*_*b*_(*χ*_1_). To begin, however, the user will first redefine the vector **s** whose components are now the *d* order parameters considered useful in constructing this new local linear RC component. The purpose of this redefinition is to capture a direction not expressible through the original *k*-dimensional subspace from which we optimized *χ*_1_ but that describes the local behavior of the slow degree of freedom. The procedure for choosing the *d* new order parameters will be as usual context specific. For ligand-protein unbinding, which is the interest of this publication, these are mostly the distances between the ligand center-of-mass and protein residues. Multi-RAVE now launches an MD simulation that is biased along *χ*_1_ using the current converged estimate of *V*_*b*_(*χ*_1_) and tracks the time evolution of the redefined order parameter vector **s**.

**Figure 2:**
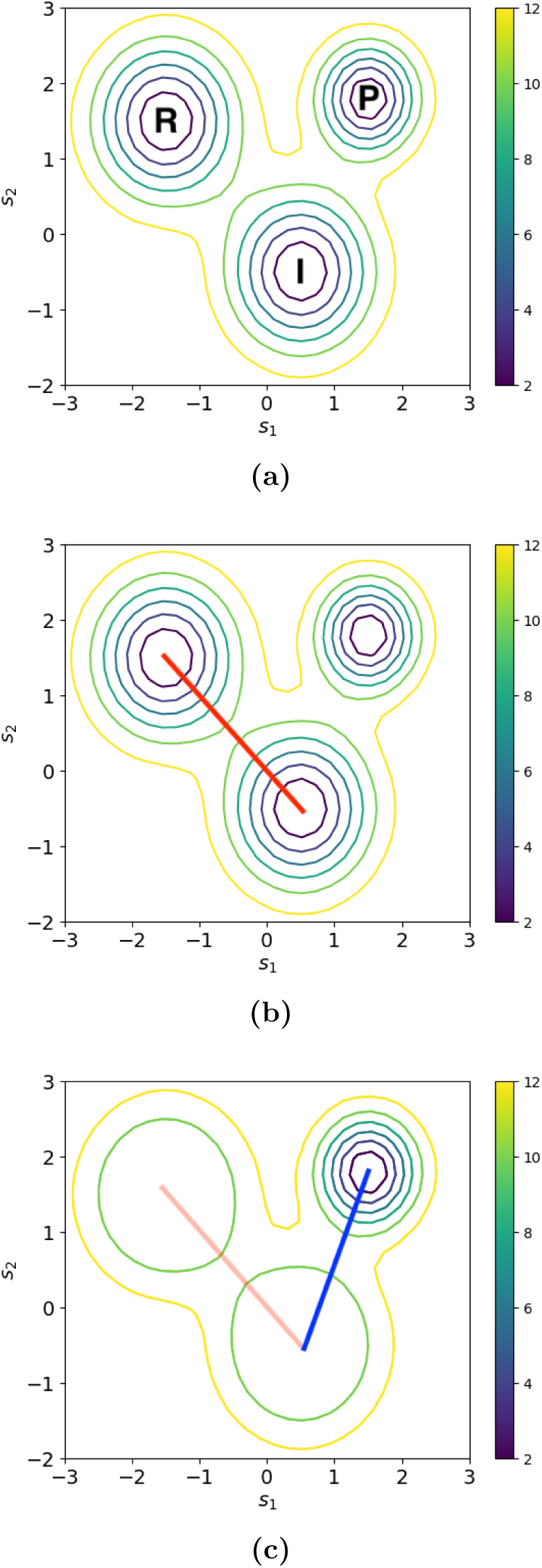
(a)A three-centered model potential with labels for the reactant (R), intermediate (I) and product (P) states. (b)Movement from left to right between the reactant to the product via an intermediate state cannot be captured with a single linear RC. (c)The three-centered model potential when the bias acting along the first RC component is not accounted for with the reweighting formula given in Eq. 4.

The multi-RAVE protocol for optimizing the second set of biasing parameters *χ*_2_ and *V*_*b*_(*χ*_2_) is similar to performing the original RAVE protocol but with a simple, essential difference that we label as the “washing out trick”. It is our intention to learn, through *χ*_2_, the features that have not already been captured by *χ*_1_. We thus need a computationally easy way to turn off the features that we have already captured. Multi-RAVE implements this task in an automated manner by optimizing *χ*_2_ without reweighting for the effect of the bias *V*_*b*_(*χ*_1_) along *χ*_1_, even though that sampling was performed using a bias. Effectively this amounts to tempering the probability distribution along *χ*_1_ so that any associated features are washed out. The net effect of this procedure is that the sampling incorporated into *χ*_1_ appears featureless so that, during the optimization of *χ*_2_ and *V*_*b*_(*χ*_2_), there is nothing to be learned from previously explored configuration space regions.

Let us be more concrete about the protocol for learning *χ*_2_ and *V*_*b*_(*χ*_2_). As we have mentioned multi-RAVE generates a time series (**s**^1^, **s**^2^, *…***s**^*n*^) using the converged estimate of *V*_*b*_(*χ*_1_). With the VAE we extract *z* and *P* (*z*) that will be our benchmark for screening through *χ*_2_ = **c s** = *c*_1_*s*_1_ + *c*_2_*s*_2_ + *…* + *c*_*d*_*s*_*d*_and corresponding *P* (*χ*_2_) via the KL divergence metric in Eq. (1). If we wished to ensure that the probabilities were unbiased we would apply the reweighting formula:

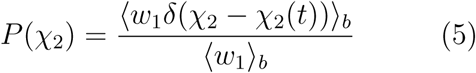

where in Eq. (5) 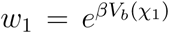. Multi-RAVE simply sets *w*_1_ = 1 to turn off the reweighting and effectively wash out any features learnt so far (see Fig. 2a vs. 2c). We thus arrive at a component of the RC *χ*_2_ and its probability distribution *P* (*χ*_2_) capturing the features of the configuration space previously uncaptured by *χ*_1_. These directly give us a second set of biasing parameters *χ*_2_ and *V*_*b*_(*χ*_2_) that are optimum given the quality of the sampling so far. We continue by iterating between the rounds of MD and VAE as per the RAVE protocol making sure to apply the washing out trick to the first set of biasing parameters *χ*_1_ and *V*_*b*_(*χ*_1_) but not the second, until the second RC and associated bias also converge. Without loss of generality, the same treatment then applies to all additional components *χ*_*i*_ and their associated biases *V*_*b*_(*χ*_*i*_) for *i* ≥ 2. We proceed to introduce local components until their converged biasing parameters lead to sampling the full rare event. Until this is accomplished, the protocol is to simply for a given RC-component number *i*, ignore all biases for components 1 to *i*-1 by setting corresponding weights *w*_*i*_ to 1 while optimizing the *i*th component and bias.

### 2.3 Binding Free Energy Calculation

While the protocol we have described here applies to generic systems, our focus in this work is on the calculation of absolute binding free energies of ligand-protein complexes, which is a challenging and important problem.^25,30^ Without loss of generality, we give illustrative examples for the benzene-T4L complex studied in this work and described in detail in Sec. 3. When the multi-RAVE protocol has converged in its estimate of a piecewise linear RC and associated biases, we launch independent MD simulations from the bound pose of the ligand-protein complex using these optimal biasing parameters. Unlike other autoencoder based enhanced sampling work, no additional biasing as in metadynamics or umbrella sampling is required.^12,15^ For the benzene-T4L complex studied here, the learnt piecewise RC and associated bias led to unbinding in simulation times between 3 to 50 nanoseconds when the natural process occurs on 100 ms or slower timescales,^31,33–35^ which is reflective of the caliber of the biasing parameters that multi-RAVE has generated.

With 20 out of 20 MD simulations sampling the benzene-T4L unbinding rare event, ligand-protein binding free energy, Δ*G*_*b*_, can then be directly calculated from the free energy profiles estimated from these runs once these have converged. In order to calculate the binding free energy we have used the potential of mean force (PMF) along the distance between the benzene center-of-mass and Tyr88. At the benzene-T4L bound position benzene is proximal to Tyr88. We then defined the reactant state along this PMF to include all values less than 0.8 *Å* and the product state to be all values greater than 1.8 *Å*. We include a standard correction term^30,33^ that accounts for the different concentration of benzene used in experiments versus simulations (1 M vs. 5 mM in the present work). One crucial caveat needs to be addressed for calculation of binding free energies from simulations such as ours once the ligand leaves the protein, it is free to explore the full solvent making it extremely rare for it to bind back. In principle, we could train RAVE to address this entropic portion of the configuration space as well. However, here for computational ease, we apply a soft restraining potential that brings the ligand back into the protein once it unbinds. The effect of this restraining potential is reweighted out in the calculation of Δ*G*_*b*_. See Supplemental Information (SI) for further details of this and other calculations in this work.

### 2.4 Simulation and Neural Network Details

The MD simulations in this work have been performed using the software GROMACS version 5.0^46^ patched with PLUMED version 2.3. ^47^ These simulations were all performed in the constant number, pressure and temperature (NPT) ensemble, with ∼10,000 water molecules in a periodic box where all side lengths are seven nanometers. The pressure of the simulation was kept at 1.0 bar and the temperature at 298 K. Constant pressure was maintained using Parrinello-Rahman barostat^48^ while the temperature was maintained constant with the v-rescale thermostat.^49^ In addition, the CHARMM22* force-field^50^ was used to describe the system. All simulations were run for ∼300 picoseconds initially, although in later RAVE rounds the MD simulation times were increased to ∼2 ns (see Sec. 3.2 for additional information).

The current VAE implementation uses a neural network architecture almost identical to the set-up used in the original RAVE publication.^18^ The input and reconstruction spaces are 2dimensional (i.e. 2-dimensional order parameter vectors) while the probabilistic encoder and decoder are each 3 layers of 512-dimensional vectors. The latent variable representation of the RC is 1-dimensional. The final layer of the encoder did not have an activation function, meaning it was defined as a linear transformation. The output layer of the decoder used tanh as the activation function. We refer the readers to the original RAVE publication for a schematic illustration of the VAE architecture as well as additional details regarding the architecture.^18^

In order to implement the aforementioned VAE architecture and to train it we have used the high level deep learning library called Keras.^51^ The training of the neural network was performed using the RMSprop algorithm, which is a variation of stochastic gradient descent. The learning rate used with RMSprop was 0.0001 or 0.0002 depending on the size of the dataset. Training was performed over 1000-3000 epochs also depending on the size of the dataset. In total, 34 rounds of MD–VAE were used to train the RC and the time-independent static biases along them.

## 3. RESULTS

### 3.1 Binding Free Energy

We have performed in total 20 biased MD simulations, all of them initiated from the benzeneT4L bound configuration using the RC and bias identified from construction rounds of RAVE. 6 of these trajectories ran for longer than 100 ns in order to monitor the detailed convergence of the Δ*G*_*b*_ estimate. As can be seen in Fig. 4, this estimate appears to be converging before the simulation time reaches 100 ns. We thus obtain, using 100 ns, a Δ*G*_*b*_ estimate of -8.2 ±0.8 kcal/mol, although similar results would be obtained with much a shorter simulation time (see the behavior of the time-dependent average given as the black curve in Fig. 4). This result is in good agreement with other calculations reported in the literature, for example -6.9 ± 0.8 kcal/mol from Mondal et al. ^34^ based on long unbiased simulations and subsequent application of Markov State Modeling, and -5.96 ± 0.19 kcal/mol from the alchemical-based calculations of Deng and Roux ^25^. Notice in Fig. 4 that all 6 trajectories underestimate these reported values suggesting that our Δ*G*_*b*_ estimates might be a lower bound. It is possible that this arises from the slower rate of sampling the unbound state as opposed to the initial metastable well corresponding to the bound complex, as we have used a sub-optimal restraining potential to bring the ligand back into the protein (see SI). Whether this is true in general, however, will be the subject of future investigations.

**Figure 4:**
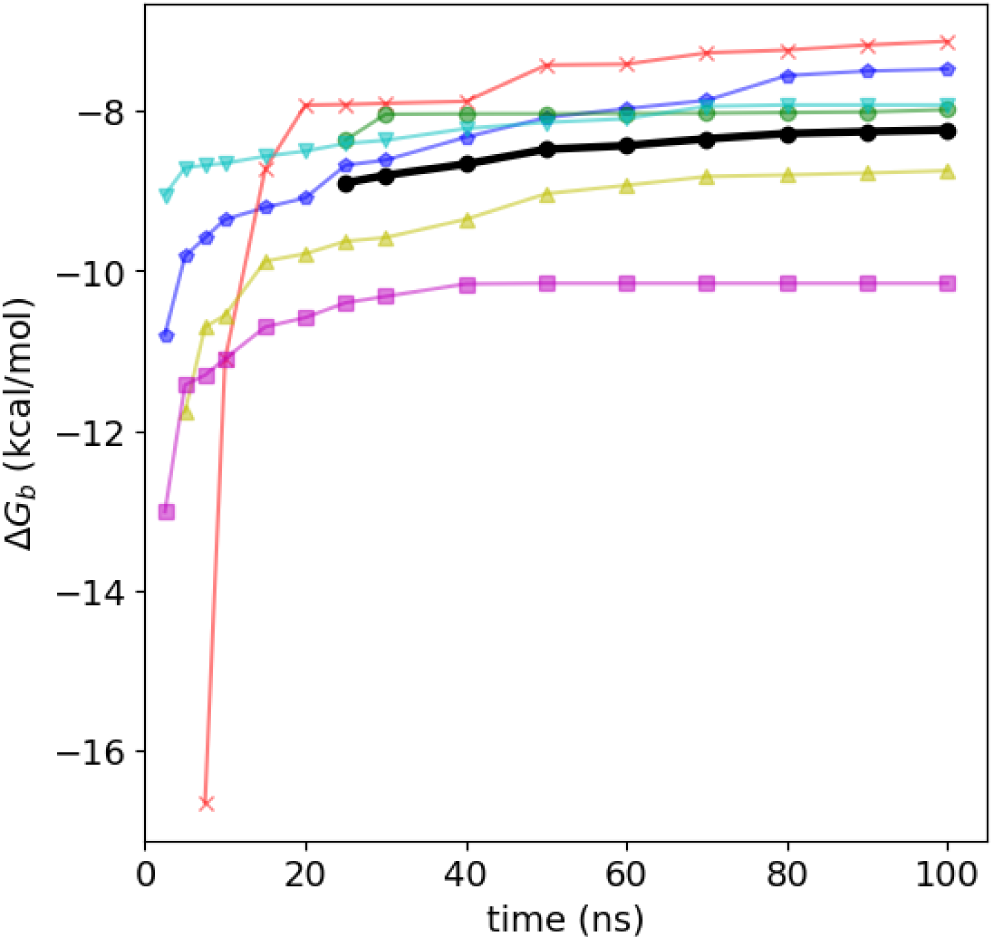
Calculated binding free energy as a function of time. Six different trajectories, in color, are shown, while the time-dependent average value is given as the black line represents.

### 3.2 Order Parameters and Reaction Coordinate

We show in Table 1 the T4L residues we used to define the distances relative to the benzene center-of-mass. Our RC components were eventually based on using just these distance-based order parameters. We also considered other order parameters such as protein inter-residue distances and protein residue hydration states, but these had close to 0 weight in the RC on a consistent basis. We will thus not describe them further here. To drive the ligand to unbind from the protein, four local linear RC components was found to be sufficient. These components were learnt sequentially using RAVE:

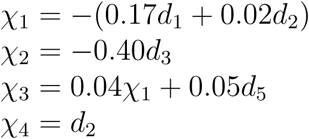

with *d*_*i*_, *i* = 1, 2, 3, 4, 5, defined as the distance between the center-of-mass of benzene and the protein residues according to Table 1. In terms of the snapshots in Figs. 3a and 3b, *d*_1_ measures the distance of benzene from helix 4, *d*_2_ from helix 5, *d*_3_ from helix 8, *d*_4_ from helix 7 and *d*_5_ from helix 6. Notice that *χ*_1_ is composed of *d*_1_ and *d*_2_, which have been defined using residues on helices that when in close contact with benzene corresponds to the initial reactant state. *χ*_2_, meanwhile, is defined in terms of *d*_3_ whose associated residue in close contact with benzene corresponds to intermediate metastable state.

**Table 1:**
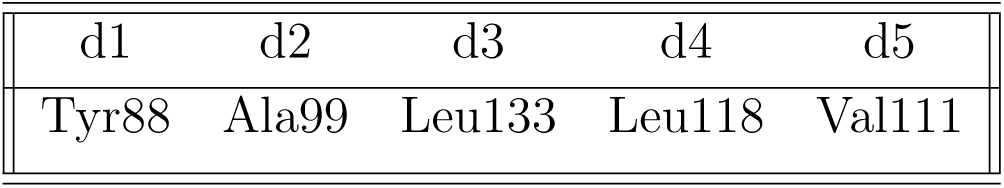
T4L protein residues used to define the order parameter distances, relative to the benzene center-of-mass, used in the construction of the piecewise linear RC.

| d1 | d2 | d3 | d4 | d5 |
| --- | --- | --- | --- | --- |
| Tyr88 | Ala99 | Leu133 | Leu118 | Val111 |

**Figure 3:**
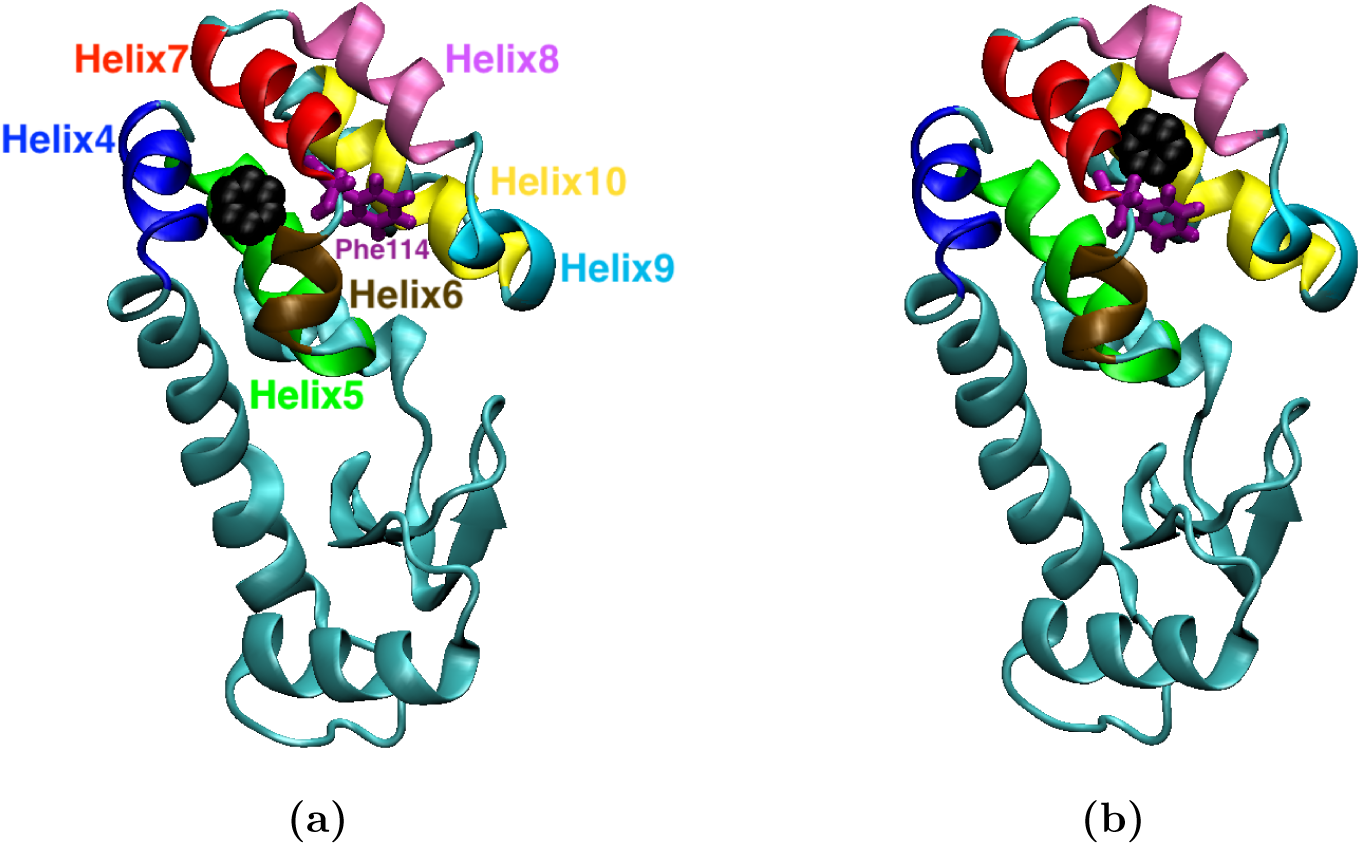
(a)Snapshot of the benzene-T4L complex in the bound state configuration from which all trajectories are initiated. (b)Snapshot of the benzene-T4L complex in the intermediate metastable state from which it escapes into the solvent through the helices 6, 7, 8 and 9.

We began learning the first of these four linear RC components using extremely short simulations of ∼300 picoseconds. These simulation times were enough to converge a first RC component but were not sufficient to learn a second RC component. For this reason we increased the simulation time for the remaining RC components to be trained to be ∼2 ns. We will investigate in future work the critical question of the simulation length which should be used to maximize the efficiency and accuracy of RAVE. Once we had learned linear RC components capable of driving exploration of configuration space from the tightly bound complex into higher energy regions and eventually into escape into the solvent we seized to continue training. Hence we did not attempt to train a RC and bias for capturing re-entry into the protein, and instead used simple restraining potentials to bring the ligand back.

As mentioned in Sec. 3.1, we ran 20 independent biased MD simulations, all of which led to benzene unbinding from the T4L buried binding site. Often, the unbinding event occurred in the first few ns of simulation although in one case benzene unbinding was observed after 50 ns. Prior to unbinding, however, we observed that benzene tends to show back and forth movement between being proximal to either helix 4 or helix 7, as shown in Fig. 5. This hopping between wells corresponds to sampling the initial bound state and an intermediate state, which suggests that RAVE can learn biasing parameters capable of reversibly sampling the reaction path until ligand escape into the solvent occurs. Taken together, these demonstrate the quality and reliability of the RC and bias so constructed.

**Figure 5:**
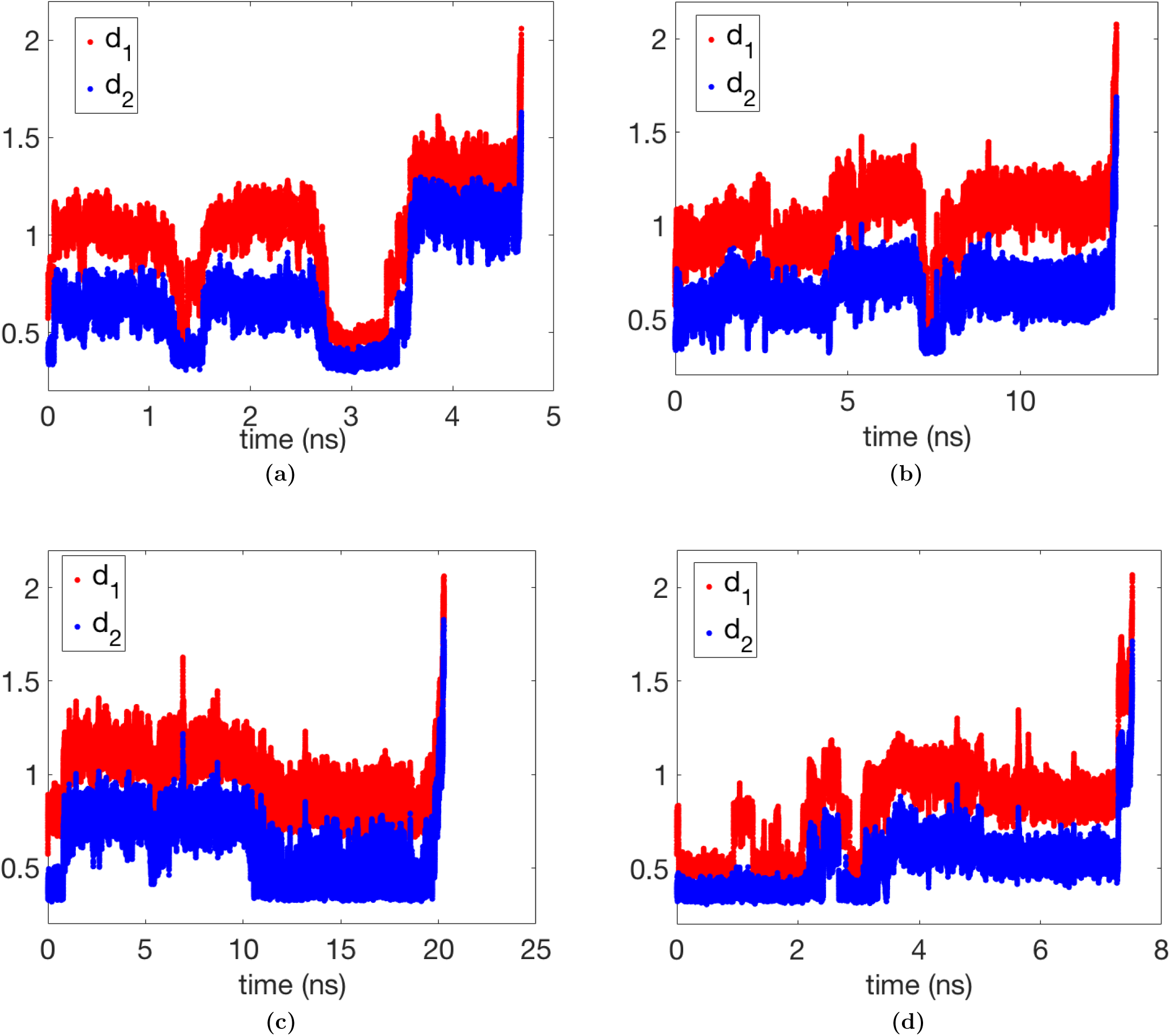
(a-d) show four (out of twenty total) independent biased trajectories in terms of order parameters *d*_1_ (red) and *d*_2_ (blue). Clear back-and-forth movement between various metastable states can be seen. Also note the short but different first passage times to unbinding for different trajectories.

### 3.3 Unbinding Mechanism

A range of specialized MD based enhanced sampling methods have been used to uncover several paths corresponding to benzene escape from the buried binding site of T4L.^31,34,35^ Many of these methods have also attempted to quantify the actual dissociation and association rate constants. Here we do not attempt to quantify these constants (which will be the subject of future work). However, our biased trajectories do show clear back and forth movement between various metastable states (See Fig. 5 showing 4 out of 20 trajectories in terms of the *d*_1_ and *d*_2_ order parameters), which is a hallmark of lack of hysteresis in enhanced sampling simulations, and is typically rather difficult to achieve.^10^ It is this easy inter-state movement which is what leads us to classify our unbinding as reversible, and gives us confidence to draw mechanistic conclusions on the basis of these trajectories. Our optimized piecewise linear RC and associated bias led to eventual benzene escape from the binding pocket and into the solvent through helices 6, 7, 8 and 9, via a metastable intermediate state corresponding to the benzene being in simultaneous close contact with helices 7 and 8. Figs. 3a and 3b show snapshots of the benzene-T4L bound pose and intermediate metastable state. The presence of these two states is also shown in Figs. 6a–6c. It is interesting to note that movement from the benzene-T4L bound position to the intermediate is correlated to slight increases in the “transient motions” of helices 7 and 8. The relevance of different short-lived protein “breathing motions” for making the deeply buried binding site accessible to ligands has been pointed by various other theoretical studies.^32,34,35,52^ Escape of the benzene into the solvent appears to require these slight increases in helix-helix distance, as shown in Fig. 6b. A movie of the unbinding simulation showing these events is provided in the SI.

**Figure 6:**
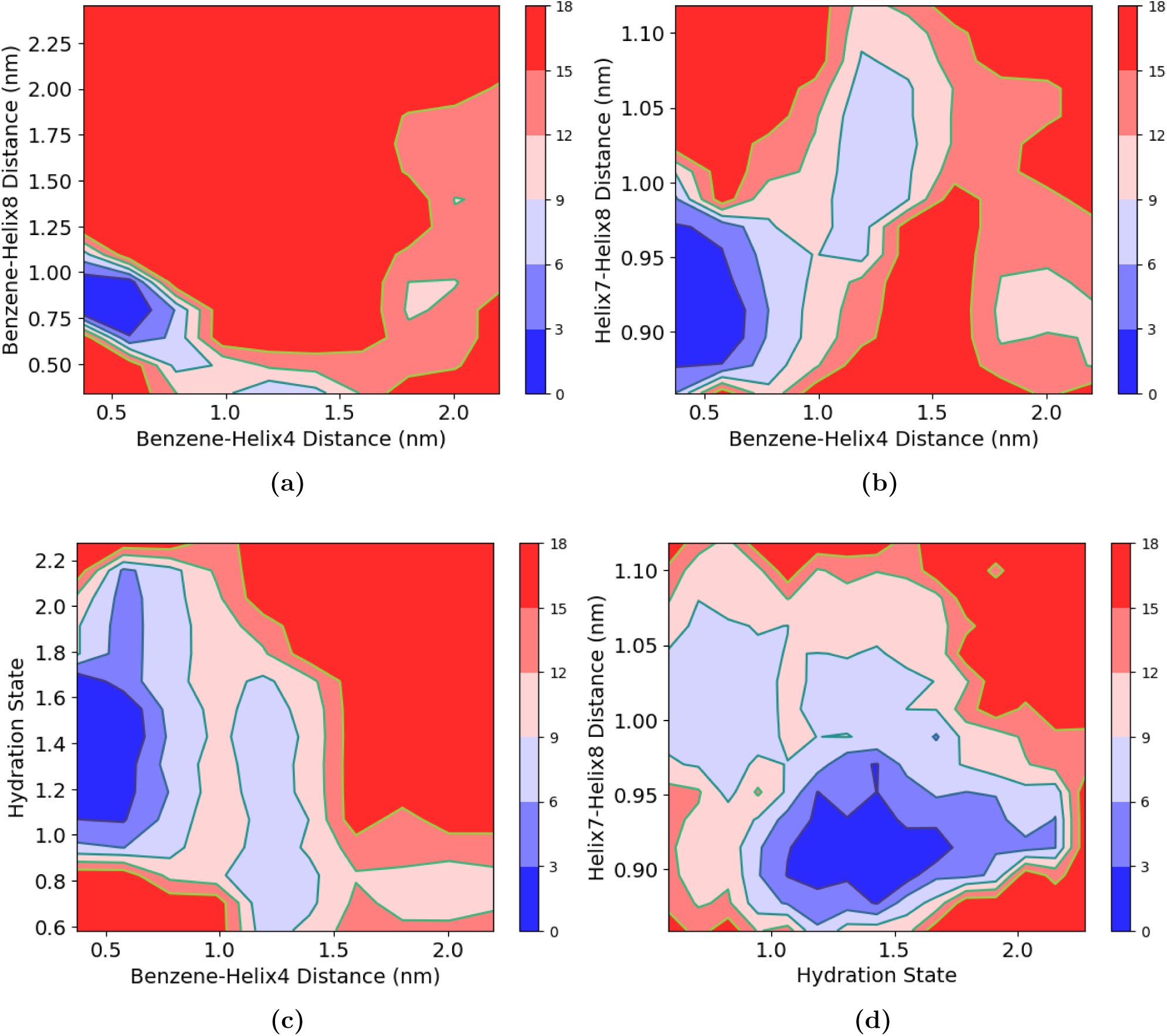
Contour plots of free energies, in kcal/mol, showing the escape pathway when projected onto different two-dimensional order parameters. The helix numbering follows the convention given in Fig. 3a. (a)Free energy profile given as a function of the main components of *χ*_1_ and *χ*_3_. (b)Free energy profile given as a function of distance from binding site and helix7-helix8 breathing motion. (c) Free energy profile given as a function of distance from binding site and hydration state between helices 8 and 10. (d)Free energy profile given as a function of hydration state between helices 8 and 10 and helix7-helix8 distance.

Interestingly, our initial benzene-T4L complex has two short-lived water molecules in the non-polar cavity between helices 8 and 10. The timescale for water exit tends to be much faster than benzene exit from the binding pocket, found to be less than 100 ns in our unbiased MD simulations. Similar waters have been reported in experiments for many variants of T4L and also from simulations. ^52^ As shown in Fig. 6c, the movement from the intermediate to the outside of the binding pocket occurs post-water exit. Sampling of the intermediate metastable state does take place while one water molecule either remains close to its initial position or enters the binding pocket before eventual escape into the solvent. The exit of water molecules from the binding pocket is thus a relatively fast, but mandatory event for ligand unbinding to occur.

## 4 DISCUSSION

Over the past decade a tremendous number of machine learning approaches relevant to various aspects of molecular simulations have become available in the literature.^12–15,53–58^ A major open problem in the field has been whether machine learning can be leveraged to make rare events less rare, therefore making amenable their accurate sampling via standard computational resources. Our recent publication marked a promising step forward in this direction.^18^ Through the use of an iterative deep learning-MD scheme we demonstrated that significant enhancement in the sampling of model potentials could be achieved. In the current work we have extended the applicability of RAVE by showing how it can be used to simultaneously learn the reaction coordinate and calculate the absolute binding free energy in a much more challenging test case, namely the benzene-T4L system in explicit water. A simple but crucial methodological extension of RAVE has been introduced here, named multi-RAVE, that allows learning a RC with multiple components and explicit dependence on location in configuration space. It is this position dependence that allows multi-RAVE to construct an overall non-linear RC as a sum of piecewise linear components that nonetheless captures the slow non-linear degree of freedom, with the added benefit of allowing for clear physical interpretability.

RAVE shares some similarities with Diffusion Map-directed MD ^59^ and intrinsic map dynamics^60^ which also perform sampling without pre-knowledge of low dimensional RC. Similar to RAVE, in these methods as well the exploration of the unknown configuration space is broken down into local components. In Diffusion Map-direction MD, for instance, localized slow diffusion coordinates are determined from an MD simulation. Intrinsic map dynamics, meanwhile, learns local parametrizations to the low-dimensional manifold that describes the slow process of interest. Both methods launch an MD simulation from the edge of the previously explored region in order to start a new iteration. RAVE is different from these methods because although it is learning local RCs from local regions of the configuration space, it is in principle independent of launching the simulation from the boundary between two regions due to the reweighting of just the “local” order parameters. That is, the final bias potential and RC learnt in RAVE allow complete movement from any point A to point B in configuration space in extra simulations with no further training or time-dependent biasing needed.

The set of RC components and bias potentials we used to describe the entire benzene-T4L unbinding process had four members (refered to in the text as 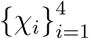 and 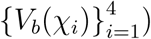. Although we were able to get full unbinding using all biases at the same time, we found that the simulation time to unbind became much smaller (less than 10ns on average from earlier around 50ns or higher) if we switched once between the two sets of biases depending on which part of the landscape the trajectory commits to. This requires a labeling of the metastable states, and thus is a form of supervised learning. In future work we will address the question of how to achieve full unbinding without having the need to switch biases on and off depending on which metastable state the trajectory had committed to. That we were able to accomplish unbinding with only one switch is not trivial this was only possible due to the quality of the RC and the bias learnt from deep learning. To use parlance from umbrella sampling, this is in a sense equivalent to sampling the full energy landscape of benzene-lyzozyme unbinding with only two umbrella potentials. No such reported work exists, to the best of our knowledge.

In the work presented here we have shown how our recent RAVE algorithm^18^ can be used to investigate the unbinding of realistic ligand-protein complexes. It is interesting to note that unbinding could be accomplished using geometric distances, without having to resort to protein-protein or solvation order parameters, whose weights were identified by RAVE to be consistently close to 0. We have noticed that the use of different distances could in principle lead to sampling different unbinding paths. Although we leave accurate kinetics of ligand-protein unbinding for upcoming work, that RAVE seems capable of reversibly sampling at least one path reversibly and in principle different paths as well seems to suggest promise for obtaining accurate kinetics, possibly through the use of an acceleration factor approach.^61–63^ We also hope to be able to apply RAVE to other established benchmarks in the field such as the SAMPL challenges for blind prediction of host–guest binding affinities.^64^

To our knowledge the current work is the first to obtain reasonable absolute Δ*G*_*b*_ estimates for the benzene-T4L complex using a PMF based approach. Wang et al. ^33^ attempted to obtain such an estimate but were not able to converge Δ*G*_*b*_ despite long simulation times. In addition, Miao et al. obtained one binding event and thus produced estimates subject to high error. ^31^ It is often thought of as less convenient to use free energy surface or potential of mean force (PMF) based approach, as opposed to alchemical-based approaches when the bound pose corresponds to a ligand buried deep within the binding pocket of a protein instead of the protein’s surface. In addition, a big criticism of PMF based approaches has also been their sensitivity to the chosen RC to perform the sampling along. RAVE addresses all of these issues – it learns a RC on-the-fly and gives accurate free energy estimates in nominal simulation time.

An open-source software implementing RAVE in its orignal form as well as the multidimensional extension introduced here, will soon be released for use by the wider community. We hope the current work makes a strong case for the use of RAVE as a method that uses deep learning to construct both the reaction coordinate and associated free energy profile in complex molecular systems, a problem that has been one of the holy grails of the field.

## 5 ACKNOWLEDGMENTS

We would like to thank both Yong Wang and Kresten Lindorff-Larsen for sharing their GROMACS input files for the L99A lysozyme-benzene complex. We would also like to thank Pablo Bravo for his essential work in helping develop the original RAVE protocol. We thank Netnaset Woldegerima for her thoughtful insights as well both Yihang Wang, Zachary Smith and Freddy Alexis Cisneros for their careful reading of this manuscript. We also thank Deepthought2, MARCC and XSEDE (projects CHE180007P and CHE180027P) for providing the computational resources used to perform this work. PT would like to thank University of Maryland Graduate School for financial support through the Research and Scholarship Award (RASA).

